# Ankyrin-B is required for the establishment and maintenance of lens cytoarchitecture, mechanics, and clarity

**DOI:** 10.1101/2024.06.12.598702

**Authors:** Rupalatha Maddala, Ariana Allen, Nikolai P. Skiba, Ponugoti Vasantha Rao

## Abstract

**Summary Statement:** This study illustrates a vital role for ankyrin-B in lens architecture, growth and function through its involvement in membrane protein and spectrin-actin cytoskeletal organization and stability

The transparent ocular lens is essential for vision by focusing light onto the retina. Despite recognizing the importance of its unique cellular architecture and mechanical properties, the molecular mechanisms governing these attributes remain elusive. This study aims to elucidate the role of ankyrin-B (AnkB), a membrane scaffolding protein, in lens cytoarchitecture, growth and function using a conditional knockout (cKO) mouse model. AnkB cKO mouse has no defects in lens morphogenesis, but exhibited changes that supported a global role for AnkB in maintenance of lens clarity, size, cytoarchitecture, and stiffness. Notably, absence of AnkB led to nuclear cataract formation, evident from P16. AnkB cKO lens fibers exhibit progressive disruption in membrane organization of the spectrin-actin cytoskeleton, channel proteins, cell-cell adhesion, shape change, loss and degradation of several membrane proteins (e.g., NrCAM. N-cadherin and aquaporin-0) along with a disorganized plasma membrane and impaired ball-and-socket membrane interdigitations. Furthermore, absence of AnkB led to decreased lens stiffness. Collectively, these results illustrate the essential role for AnkB in lens architecture, growth and function through its involvement in membrane protein and cytoskeletal organization.

## INTRODUCTION

The ocular transparent and avascular lens is indispensable for vision, precisely focusing light onto the retina. Across many species, including humans, the lens continues to grow throughout life, comprising a monolayer of epithelium covering the anterior surface and elongated, differentiated long fiber cells within a thick collagenous capsule (Quinlan and Clark, 2022, Hejtmancik et al., 2015). Numerous cellular attributes are pivotal for maintaining lens transparency and accommodation, including the differentiation of epithelial cells into ribbon-like long fiber cells, compact cell-cell adhesion, hexagonal symmetry, lateral membrane interdigitations, and organized membrane cytoskeletal, cell adhesion, channel, and transport proteins (Quinlan and Clark, 2022, Mathias et al., 2010, Bassnett et al., 2011, Taylor et al., 1996, Gu et al., 2019, Cvekl and Ashery-Padan, 2014, Cheng et al., 2017). Additionally, there is a programed degradation and elimination of most cellular organelles in lens fibers (Bassnett, 1997). While extensive research has focused on understanding the molecular mechanisms maintaining lens transparency and mechanical properties (Bassnett et al., 2011, Quinlan and Clark, 2022, Cheng et al., 2017, Hejtmancik et al., 2015, Cheng et al., 2018, Wang et al., 2016), our understanding of the role of membrane scaffolding proteins in regulating lens fiber cell geometry, adhesion, membrane organization, and mechanical properties remains limited (Maddala et al., 2016, More et al., 2001, Maddala et al., 2011).

Ankyrins, conserved metazoan scaffolding proteins, play a crucial role in tethering the spectrin-actin cytoskeleton to various membrane proteins, including channels, transporters, cell adhesion molecules, and signaling proteins, thereby regulating their membrane subdomain organization and activities (Bennett and Baines, 2001, Bennett and Lorenzo, 2016, Srinivasan et al., 1988). Mammals express three ankyrin subtypes encoded by three distinct genes, *ANK1*, *ANK2* and *ANK3*. While *ANK1* primarily expresses in erythrocytes, encoding ankyrin-R, *ANK2* and *ANK3,* expressed in various tissues including the brain, heart, eyes, and others, encode ankyrin-B (AnkB) and ankyrin-G (AnkG), respectively (Bennett and Baines, 2001, Bennett, 1979). These genes also exhibit several splice variants expressed in different tissues (Chan et al., 1993). Ankyrin proteins contain conserved functional domains crucial for their physiological activities, including the N-terminal membrane binding domain (MBD) with 24 tandem ankyrin repeats, a spectrin-binding domain in the middle, and a C-terminal regulatory domain (Bennett and Lorenzo, 2016, Michaely and Bennett, 1995). Ankyrins interact directly with β-spectrin, L1 family of cell adhesion proteins including NrCAM, dystroglycan, channels proteins, transport proteins, and certain receptors (Mohler et al., 2004b, Davis and Bennett, 1994, Mohler et al., 2004a, Ayalon et al., 2008, Michaely and Bennett, 1995, More et al., 2001, Mohler et al., 2003). Importantly, mutations and dysregulation of ankyrins are associated with various diseases and abnormalities of the heart, brain, metabolism, and other tissues, underscoring their critical roles in cellular physiology and tissue functions (Stevens and Rasband, 2021, Mohler and Bennett, 2005, Mohler et al., 2003, Lorenzo et al., 2015).

The ocular lens abundantly expresses AnkB and to a lesser extent AnkG (one-tenth levels of AnkB) (Rasiah et al., 2019). While AnkG predominantly distributes to the lens epithelium and regulates epithelial phenotype, polarity, and lens morphogenesis, growth and function (Rasiah et al., 2019), AnkB intensely localizes to lens fibers relative to epithelium. This distribution coincides with the preferential expression of the AnkB-binding transmembrane cell adhesion protein, NrCAM in lens fibers (More et al., 2001, Maddala et al., 2016). However, the precise role of AnkB in lens fiber architecture, mechanical properties, growth and function remains unclear (More et al., 2001, Maddala et al., 2016). Extensively characterized AnkB null mice survive only for less than a day after birth, while haploinsufficient (heterozygous) mice grow and breed normally (Scotland et al., 1998). Reports suggest that older AnkB heterozygous mouse lenses exhibit haziness and certain abnormalities in lens fiber cell architecture and membrane organization without affecting lens growth (Maddala et al., 2016). However, the impact of complete AnkB absence on lens growth and function is yet to be fully elucidated (More et al., 2001). To gain further insights into the role of AnkB in lens morphogenesis, growth and function, we developed an AnkB conditional deficient mouse model (AnkB cKO) using a LoxP-Cre recombination approach. Neonatal and postnatal lenses derived from AnkB cKO mice were characterized and compared with littermate AnkB LoxP control mice. This study’s findings reveal that while AnkB is not required for lens development and differentiation, its absence impair lens growth, transparency, fiber cell cytoarchitecture, membrane organization, and mechanical properties in postnatal and adult mice.

## RESULTS

### Ankyrin-B interaction with spectrin-actin skeleton and membrane-associated proteins in lens fibers

The bulk of the ocular lens consists of differentiated fiber cells, with the plasma membrane being a predominant component (Quinlan and Clark, 2022, Hejtmancik et al., 2015, Bassnett et al., 2011). Scaffolding proteins linking cytoskeletal proteins with integral membrane proteins are crucial for maintaining organ structure and function (Bennett and Lorenzo, 2016, Bennett and Baines, 2001). However, our understanding of how interactions of key membrane scaffolding proteins regulate the unique structural and functional attributes of lens fibers remains limited (Maddala et al., 2016). AnkB is abundantly expressed in the mouse lens (P30), ranking among the top 100 most abundant proteins based on RNAseq analysis (Maddala et al., 2021). Immunoblot analysis of total lysates from P16 mouse lenses revealed a predominantly expressed splice variant of AnkB with ∼250-270 kDa, also detectable in P16 mouse brain lysates (**Fig. 1A**). Immunofluorescence analysis demonstrated intense distribution of AnkB in fibers compared to the epithelium of the mouse lens (P21; **Fig.1B**). Within fibers, AnkB exhibited uniform distribution across the long and short arms of hexagonal fiber cell membranes in the cortical region of P21 mouse lenses (**Fig. 1C**, equatorial plane).

**Figure 1.**
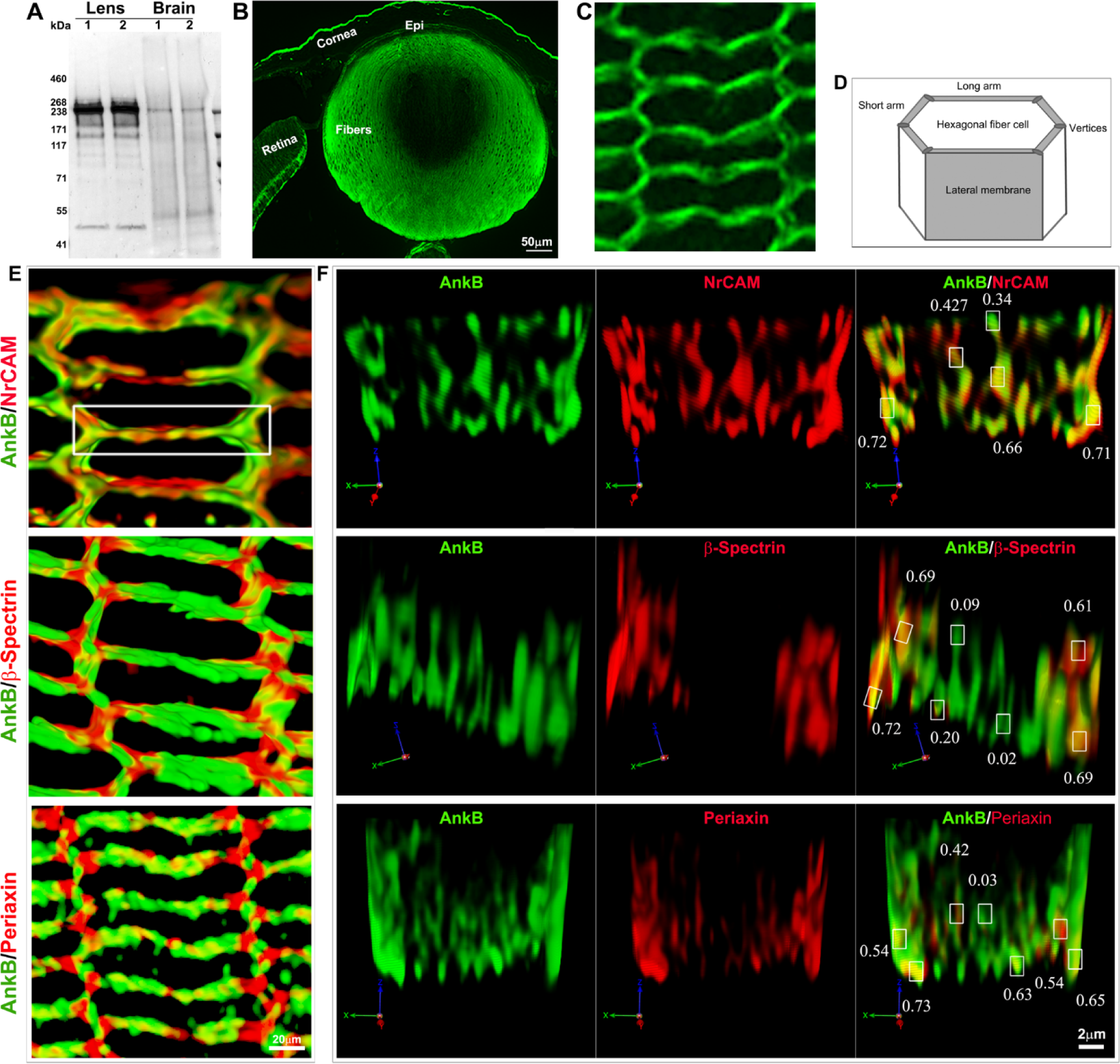
Ankyrin-B co-distribution with NrCAM, *β*-spectrin and periaxin in mouse lens fiber cell membrane subdomains. A. Immunoblot analysis of mouse lens (P16) homogenates (20 µg) demonstrates the presence of a prominent ∼250-270 kDa AnkB isoform, along with weaker molecular mass isoform or derivatives of the 250/270 kDa protein. A similar molecular mass AnkB was readily detectable in brain homogenates (2 µg). 1 & 2 are two independent samples. B & C. Immunofluorescence reveals prominent distribution of AnkB in P21 mouse lens fibers compare to the epithelium (sagittal plane), and in the hexagonal fiber cell short and long arms (equatorial plane), respectively. D. Schematic representation of the lateral membrane view of lens hexagonal fiber cell long and short arms. E & F. AnkB co-distribution with NrCAM, β-spectrin and periaxin in mouse lens hexagonal cortical fibers. En face view of immunofluorescence images (captured using Zeiss LSM 780 inverted Airyscan) of the indicated proteins in fiber cell (equatorial plane, P21 lens; E) and lateral membrane subdomains (F), analyzed using 3D deconvolution with Volocity image analysis software, depicts AnkB and NrCAM colocalization across all membrane subdomains (including long and short arms and vertices), as indicated by Pearson correlation coefficient values in panel F. While AnkB predominantly colocalizes with β-spectrin at vertices of short arms but not long arms, it shows colocalization with periaxin in specific lateral membrane subdomains at long arms and close interaction at vertices. Scale bars represent image magnification. Images are representative of analyses from 3 to 4 independent samples.

To identify AnkB’s interacting proteins in the lens, we performed immunoprecipitation analysis using an AnkB monoclonal antibody and tissue lysates (800xg supernatants) from differentiated lenses (pooled from P21 and P30 mice) alongside an IgG control. SDS-PAGE analysis of AnkB immunoprecipitates identified several proteins (Supplemental **Fig. S1A**), subsequently subjected to trypsin digestion and mass spectrometry analysis. Multiple cytoskeletal and cell adhesion proteins were identified alongside AnkB in two independent samples. Supplemental **Table S1** lists various proteins immunoprecipitated from lens lysates with the AnkB antibody, including AnkB itself, NrCAM, N-cadherin, spectrins (both α and β), actin, various cell adhesion, and certain crystallin proteins. Immunoblot analyses confirmed the presence of the mass spectrometer-identified proteins in AnkB antibody immunoprecipitates (Supplemental **Fig. S1B**). Furthermore, we assessed colocalization using Coste’s Pearson Correlation coefficient analysis. In the lens (P21) cortical region (using paraffin embedded sections, Zeiss LSM 780 inverted Airyscan confocal microscopy and Volocity software) NrCAM, β-Spectrin, Periaxin, Aquaporin-0, and β-actin exhibited coefficients of 0.854, 0.679, 0.603, 0.531 and 0.456 respectively, indicating close colocalization with AnkB in lens fibers. It is possible that some of the co-immunoprecipitated proteins with AnkB could be through non-specific interactions including crystallins which are expressed abundantly in lens fibers.

We also analyzed AnkB colocalization at lens cortical region hexagonal fiber cell lateral membrane subdomains with NrCAM, β-spectrin, and periaxin using 3D deconvolution method with Volocity image analysis software. **Fig. 1D depicts** the schematic drawing of hexagonal lens fiber cell. En face view of fiber cells in the equatorial plane (**Fig. 1E**) and lateral membrane (**Fig. 1F**) revealed extensive colocalization of AnkB and NrCAM across membrane subdomains (at long and short arms and vertices; based on the indicated Pearson correlation coefficient values), while with β-spectrin, colocalization primarily occurred at vertices of short arms but not along long arms. Conversely, with periaxin, colocalization was observed at certain lateral membrane subdomains of long arms but exhibited close interaction at vertices. Additionally, 3D rendering movies were used to determine AnkB co-distribution with β-dystroglycan, connexin-50, β-actin and β-spectrin, and noted only discrete regions of interaction, particularly between AnkB and connexin-50 (**Supplemental movies**). In summary, these protein interactions and colocalization analyses reveal AnkB’s interaction and colocalization with several proteins in lens fibers, exhibiting varying degree of interaction and relatively close association with NrCAM at multiple membrane subdomains.

### Ankyrin-B absence impairs lens growth and transparency

AnkB null mice, surviving only half a day after birth (Smith et al., 2015, Scotland et al., 1998), prompted the development of AnkB cKO mice. These were generated by crossing AnkB floxed mice, which contain loxP sites flanking exon 24 of the *ANK2* gene (**Fig. 2A**; (Smith et al., 2015) and maintained on a C57BL/6J background with lens-specific Cre recombinase expressing transgenic mice (Le-Cre mice, **Fig. 2**) as we described earlier (Maddala et al., 2015). The Le-Cre transgenic mice used in this study express Cre recombinase at embryonic day 8.75 under the control of a Pax6 P0 enhancer/promoter, with Cre being expressed in lens epithelium and fiber cells as well as other surface ectoderm-derived eye structures (Ashery-Padan et al., 2000). For comparison with AnkB cKO mice, littermate AnkB floxed controls negative for the Cre transgene were utilized. Genotyping, immunoblot and immunofluorescence analyses confirmed the conditional deletion of AnkB gene and deficiency of AnkB protein in lenses derived from AnkB cKO mice (**Fig. 2B-F**). Postnatal AnkB cKO mice (derived from F10 and higher generations), starting from postnatal day 16 (P16), exhibited bilateral cataract with intense nuclear opacification compared to littermate controls (**Fig. 2C & D**). Interestingly, similar to observations in P1 AnkB null mice (supplemental **Fig. S2**; mice were obtained from the Bennett laboratory)(Scotland et al., 1998), lens development and differentiation were not perturbed in P1 AnkB cKO mice (supplemental **Fig. S2**). Notably, lenses derived from neonatal P10 and P14 AnkB cKO mice were transparent, with normal growth and weight similar to littermate control lenses. However, beginning at P16, lenses from AnkB cKO mice showed intense opacity, especially in the nucleus (**Fig. 2G**). This phenotype progressed rapidly by P18 and P21, accompanied by a dramatic and significant decrease in lens size and weight compared to littermate AnkB floxed control lenses (**Fig. 2G**). The lens weights were significantly decreased by ∼ 11%, 58% and 85% in P16, P18 and P21 AnkB cKO mice, respectively, compared to their respective littermate controls as shown in **Fig. 2G & H**. To investigate whether this marked decrease in lens size and weight was associated with altered cell survival, lenses from P10, P14 and P16 (sagittal, cryosections) AnkB cKO mice were evaluated for apoptosis using TUNEL labelling (Supplemental **Fig. S3**). Interestingly, this analysis did not reveal increased cell death (based on Tunnel positive cells) due to apoptosis, indicating potential involvement of alternative causes (e.g., epithelial proliferation) other than apoptosis for the decreased growth of lenses in the absence of AnkB (**Fig. S3**). Additionally, a significant increase in lens weight was noted in control mice from P14 to P21, with more than a 4-fold increase, while there was only approximately a 4% increase between P10 and P14 (**Figs. 2 & S4**). In contrast, under AnkB absence, lenses exhibited a dramatic phenotype of cataract and decreased weight starting from P16 (**Figs. 2 & S4**).

**Figure 2.**
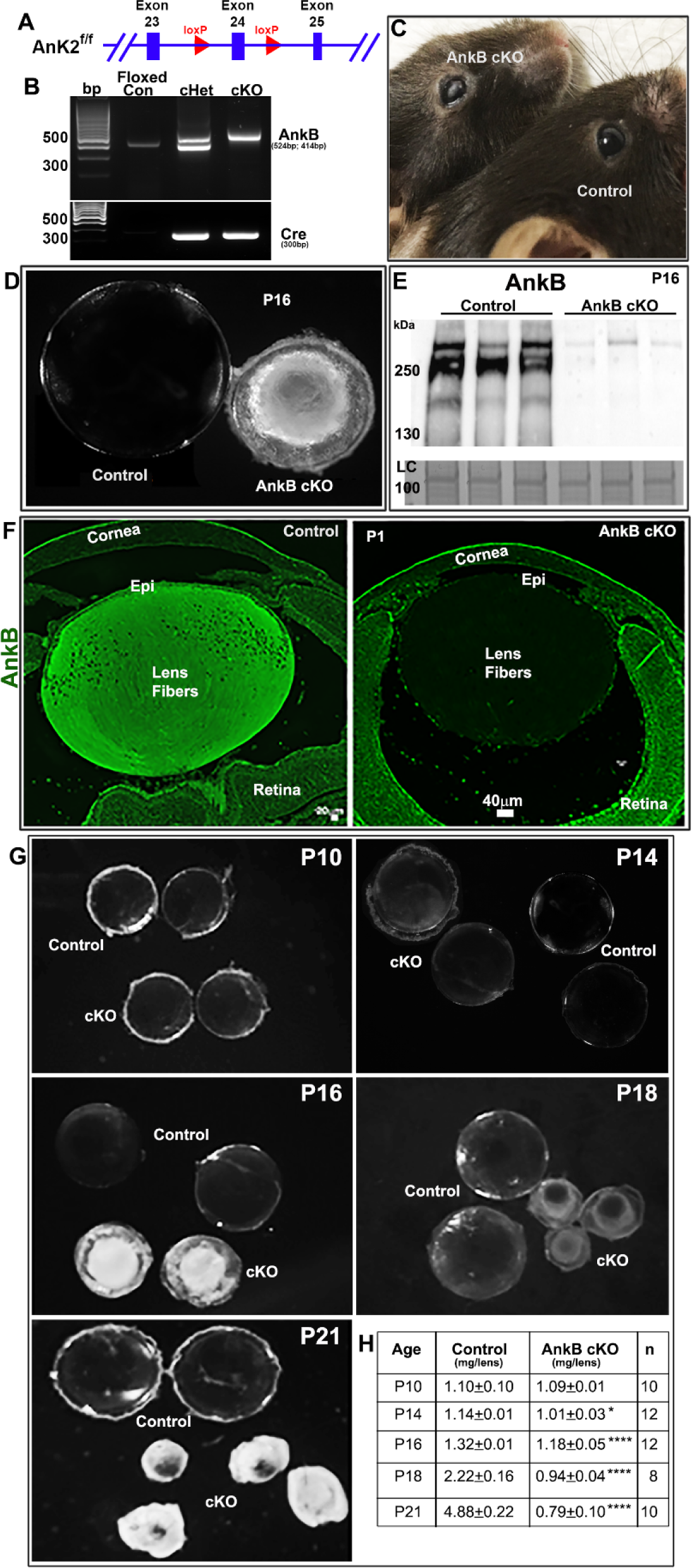
Impaired lens growth and development of nuclear cataract in AnkB cKO mice. A. AnkB cKO mice were generated by mating AnkB floxed mice (with loxP sites flanking exon 24 of the *Ank2* gene, maintained on a C57BL/6J background) with transgenic mice expressing Cre recombinase under the control of a lens-specific promoter (Le-Cre mice). B. Confirmation of AnkB cKO mouse generation through PCR analysis of tail DNA for genotyping. C. Ocular phenotype (cataract) observed in P16 AnkB cKO mice compared to littermate AnkB floxed control mice. D. P16 AnkB ckO mice exhibit markedly reduced lens size and the development of intense nuclear cataracts compared to littermate control (AnkB floxed) mice. E. Immunoblot analysis confirms the deficiency of AnkB protein in P16 AnkB cKO mouse lenses compared to control mouse lenses. F. Immunofluorescence analysis demonstrates the lens-specific deficiency of AnkB in P1 AnkB cKO mice compared to littermate control (AnkB floxed) mice. G & H. While lenses from P10 and P14 AnkB cKO mice appear transparent with minimal differences in weight and size compared to control mice, lenses from P16, P18 and P21 AnkB cKO mice show intense opacity and a significant decrease in weight compared to control mice. Epi: Epithelium, n= number of lens samples analyzed. Bar: Magnification scale. * P<0.05, **** P<0.0001.

### Ankyrin-B deficiency disrupts the lens fiber cell integrity, shape, compaction, and lateral membrane ball-and-socket interlocking digitations

Histological examination of lenses (sagittal plane cryosections with confocal microscope imaging) from P10, P14, P16, P18 and P21AnkB cKO mice, along with respective littermate controls (AnkB floxed), revealed interesting findings. Up to the P16 stage, lenses from P10 and P14 AnkB cKO mice exhibited certain sutural abnormalities but maintained fiber cell elongation, migration, and compaction compared to control lenses (**Fig. 3**). However, starting from P16, there was notable disruption in fiber organization, particularly in the lens nucleus, progressing dramatically by P18 and P21 in AnkB cKO lenses compared to controls (**Fig. 3**). This disruption extended from the nucleus to the outer cortical region, albeit without rupturing the capsule, indicating intact capsule (**Fig.3**). These histological changes correlated with decreased lens size and weight, as well as opacification (**Fig. 2**). Additionally, from P14 onwards, immunostaining of E-cadherin in the lens epithelium of AnkB CKO mice showed reduced expression compared to controls, accompanied by increased α-smooth muscle actin expression (Supplemental **Fig. S5**).

**Figure 3.**
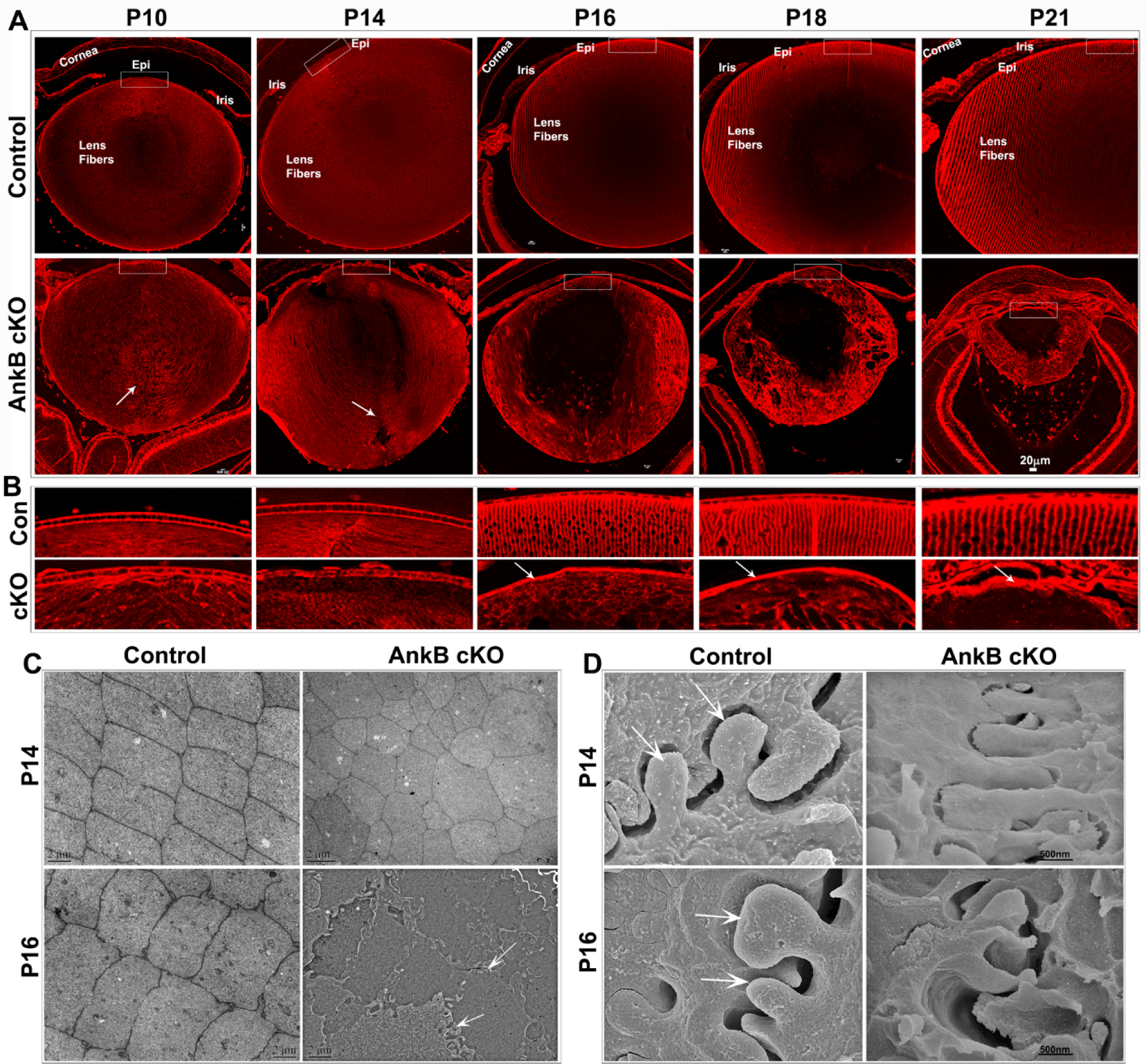
Ankyrin-B deficiency disrupts lens fiber cell morphology, organization, and compaction. A. Confocal images (single optical plane) of sagittal cryosections stained with rhodamine-phalloidin from AnkB cKO and littermate control (AnkB floxed) mice at postnatal day 10 (P10) and P14 reveal no overt abnormalities in fiber cell elongation and differentiation in AnkB cKO lenses, with some sutural abnormalities evident (arrows). However, starting from P16, notable abnormalities in fiber cell organization and shape emerge, particularly in the lens nuclear region, intensifying by P18 and P21 in AnkB cKO lenses compared to controls. Despite extensive abnormalities, the lenses of P18 and P21 AnkB cKO mice were intact. B. Interestingly, the lens epithelium in AnkB cKO mice shows fibrotic changes (arrows) starting from P16, contrasting with control lenses. C. Transmission electron microscopy analysis of P14 and P16 lenses (equatorial plane; cortical region) reveal altered fiber cell shape and organization in AnkB cKO lenses compared to uniform cuboidal shaped fibers with radial arrangement in controls. By P16, AnkB cKO lenses exhibit not only abnormal fiber shape but also extensive membrane folding (arrows) and breaks with vesicle accumulation, contrasting with the symmetric and compact organization of control fibers. D. Scanning electron microscopy of P14 and P16 sagittal sections (cortical region) from AnkB cKO mice shows impaired establishment of ball- and-socket lateral membrane interdigitations between adjacent fiber cells compared to controls (arrows). By P16, AnkB cKO lenses display rudimentary and deformed membrane formations, completely lacking ball-and-socket interdigitations, indicating a significant disruption in their establishment and maintenance under AnkB deficiency. Representative images from four independent lens samples are shown. Epi: Epithelium, Bars: Magnification scale.

Further analysis using transmission electron microscopy (TEM; Jeol 1400) revealed disrupted fiber cell shape and organization in the inner cortical region of P14 and P16 AnkB cKO lenses (equatorial plane, cortical region) compared to controls (**Fig. 3C**). Fiber cells in AnkB cKO lenses exhibited disorganization, altered shape, and extensive membrane degradation and folding (**Fig. 3C**, arrows), highlighting the role of AnkB in maintaining fiber cell shape, adhesion, organization, and membrane integrity.

Scanning electron microscope (SEM; FEI Sirion XL30-FEG) analysis of P14 and P16 AnkB cKO lenses and controls (sagittal sections; cortical region) showed differences in lateral membrane ball-and socket interdigitations between adjacent fiber cells (**Fig. 3D**). While control lenses exhibited well-developed interdigitations (**Fig. 3D**, arrows), AnkB cKO lenses displayed tube-like shapes lacking defined neck and head morphology, with a progressive deformation and loss of interdigitations by P16 (**Fig. 3D**).

Furthermore, light sheet microscopy imaging of P16 AnkB cKO lenses immunostained for aquaporin-0 revealed extensive disruptions in fiber cell compaction relative to controls (Supplemental **Fig. S6**), highlighting AnkB’s role in maintaining fiber cell architecture, adhesion, and membrane organization.

In summary, our analyses using light microscopy, TEM, SEM, and light sheet microscopy underscore the crucial role of AnkB in the morphological architecture, adhesion, and membrane organization of lens fiber cells, with AnkB deficiency significantly impairing these cellular characteristics.

### Ankyrin-B plays a crucial role in maintaining the stability of lens fiber membrane proteins

In AnkB cKO mouse lenses, there is a rapid onset of loss and degradation of membrane proteins starting from postnatal day 16 (P16). Analysis of lens soluble fractions (16000xg supernatant) at P12, P14, and P16 revealed a normal and comparable protein profile between littermate controls (AnkB floxed) and AnkB cKO mice, as observed through SDS-PAGE analysis and Coomassie blue staining (Supplemental **Fig. S7**). However, by P18, there is a noticeable decrease in the levels of soluble proteins in AnkB cKO lenses, evident from the protein band intensity (**Fig. S7**). In contrast, analysis of the membrane fraction from AnkB cKO lenses at P12, P14, and P16 displayed a decrease in the number of protein bands, as indicated by Coomassie staining, beginning at P16 and further declining dramatically by P18 compared to their respective control lenses (**Fig. 4A & B**).

**Figure. 4.**
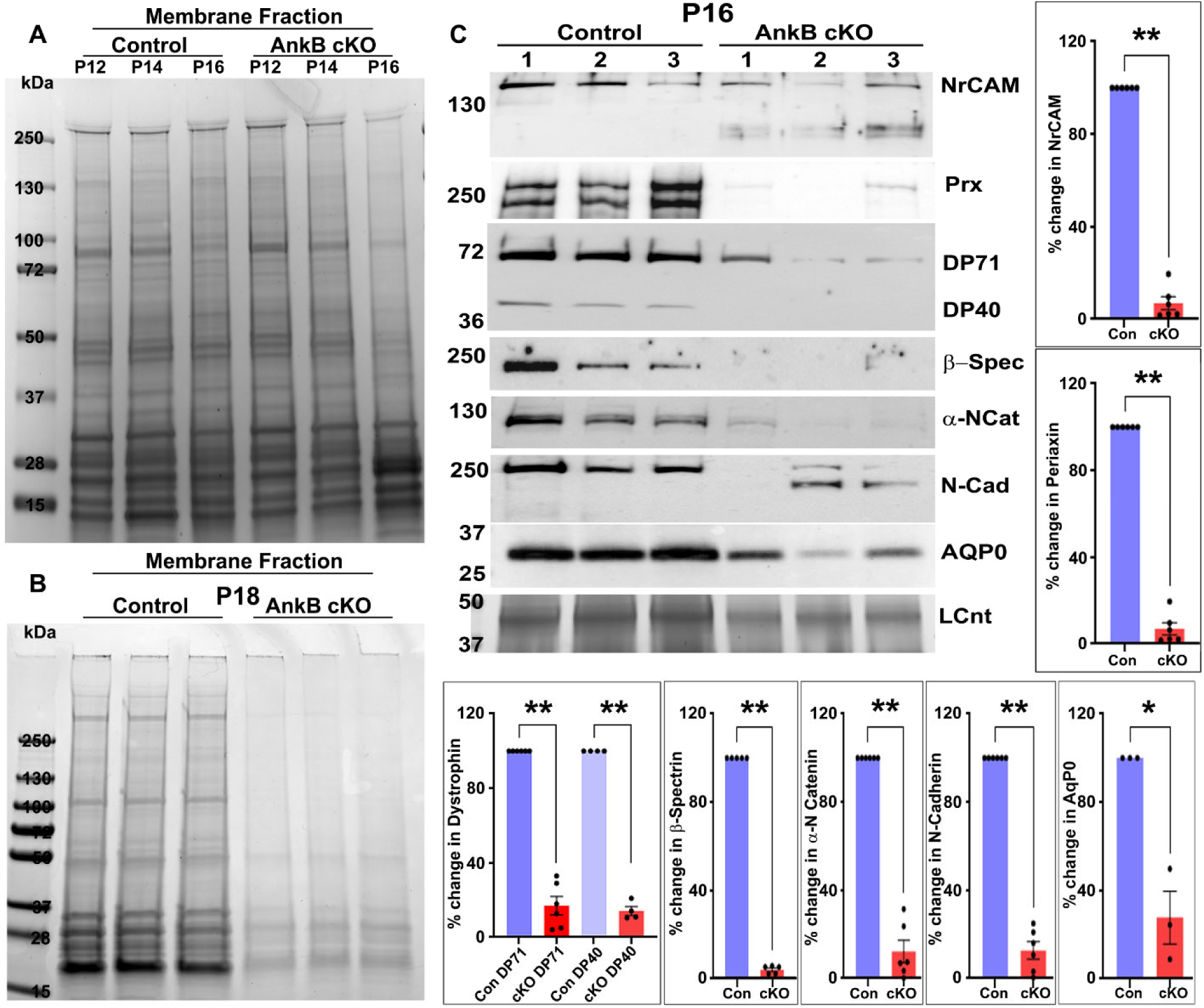
Disruption of lens fiber cell membrane protein stability under AnkB absence. A and B. SDS-PAGE separation followed by Coomassie blue staining of membrane fraction proteins extracted from AnkB cKO mouse lenses reveal a progressive disappearance and degradation of specific proteins, beginning at postnatal day 16 (P16) and markedly intensifying by P18 in comparison to their respective control (AnkB Floxed) samples, with equal amounts of protein analyzed in each case. C. Immunoblot analysis of membrane fraction proteins from P16 AnkB cKO mouse lenses demonstrates both degradation and significant reduction in the levels of NrCAM, periaxin (PRX), dystrophin (DP71 & 40), N-cadherin (N-cad), αN-catenin (αNCat), aquaporin-0 (AQP0) and β-spectrin (β-spec), as quantitated in the accompanying histograms, compared to control lenses. Lanes 1 to 3 represent three independent samples. * P<0.05. Con: Control. LCnt indicates the loading control, where protein equal amounts from membrane fractions separated by SDS-PAGE, stained with Coomassie blue, and a designated band was utilized for normalizing of loading.

Moreover, immunoblot analysis revealed a significant decrease or degradation in the levels of several membrane and membrane-associated proteins including NrCAM, periaxin, dystrophin (Dp71/Dp40), β-spectrin, aquaporin-0 (AQP-0), αN-catenin, and N-cadherin in the membrane fraction of P16 AnkB cKO lenses compared to control lenses (**Fig. 4C**). Loading controls for the membrane fraction protein analysis involved separating equal amounts of protein from lens membrane fraction samples (control and AnkB cKO) via SDS-PAGE and staining with Coomassie blue, with one of the indicated bands serving for the normalization of loading. These findings collectively suggest a decrease in the stability and perhaps expression of several membrane proteins in lenses under the absence of Ankyrin-B.

### Ankyrin-B absence disrupts fiber cell adhesion, lateral membrane protrusions, and membrane protein organization in the mouse lens

Equatorial sections of AnkB cKO mouse lenses at postnatal day 16 (P16; embedded in paraffin), displayed disruption in the hexagonal shape and radial organization of fiber cells compared to control lenses. This disruption was assessed through immunofluorescence staining analysis for β-spectrin, N-Cadherin, NrCAM, connexin-50, and β-dystroglycan, and evaluated via confocal imaging of single optical sections (**Fig. 5A**, images were from the cortical region of lens).

**Figure 5.**
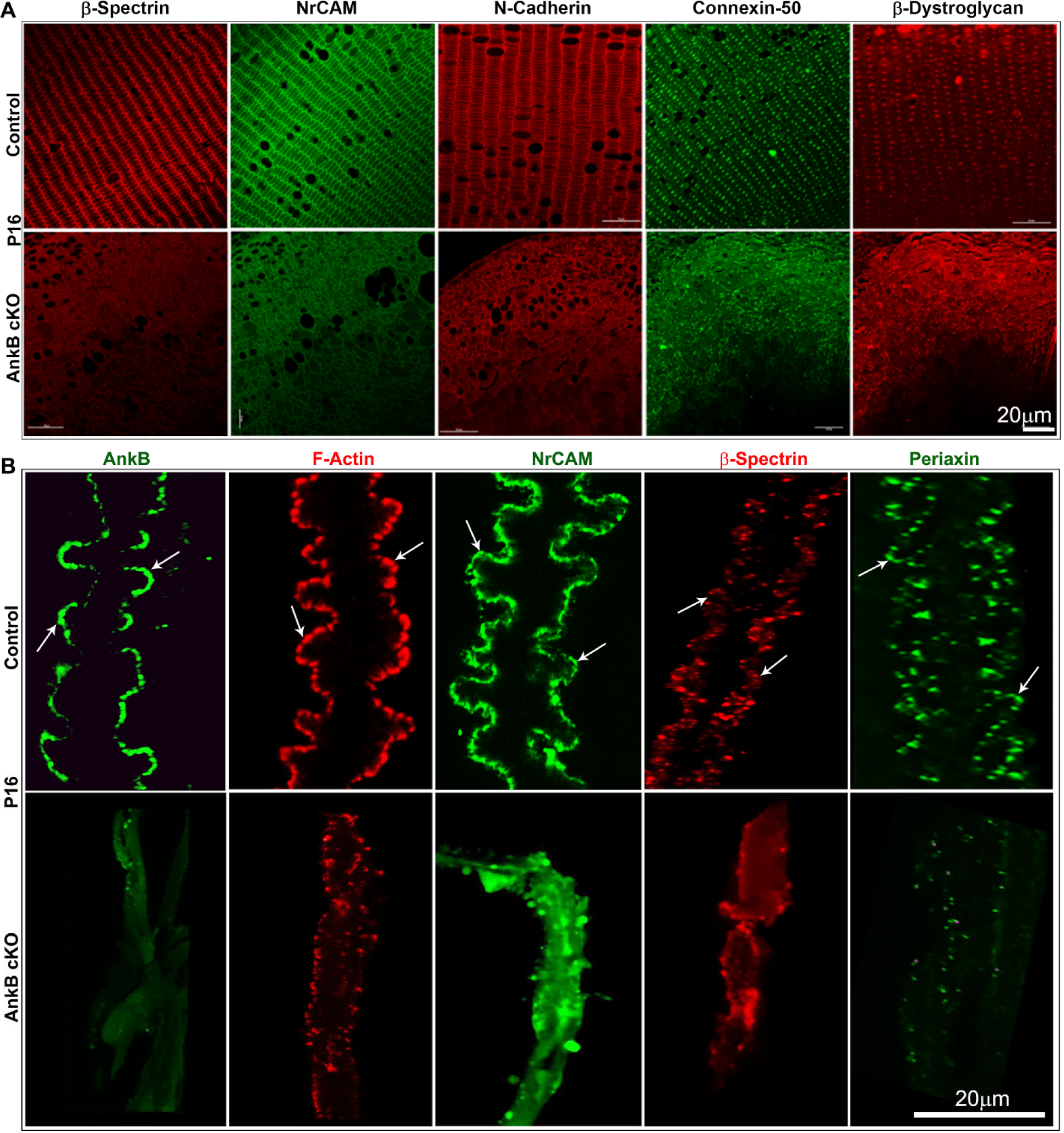
Disruption of fiber cell radial symmetry, cell adhesion, and lateral membrane protrusions in AnkB absence. A. Equatorial sections of AnkB cKO mouse lenses at P16, paraffin-embedded and immunostained for β-spectrin, NrCAM, N-Cadherin, Connexin-50 and β-dystroglycan, were imaged (single optical sections, cortical region). A significant disruption in organization and radial symmetry of fiber cells was observed compared to control (AnkB floxed) specimens. B. Single fibers peeled from the cortical region of AnkB cKO mouse lenses at P16 were shorter, thinner, and more fragile compared to fibers from control lenses. Confocal images (z stacks, maximum intensity projections) of F-actin stained (Rhodamin-phalloidine) and NrCAM, β-spectrin and periaxin immunostained single fibers from control lenses showed distinct distribution to lateral membrane protrusions and paddles (arrows). In AnkB cKO lens fibers, distribution patterns of these proteins were dramatically disrupted with no evidence of protrusions or paddles, indicating impaired organization and cell adhesive interactions. Scale bars: Magnification scale. Images are representative of analyses from 3 to 4 independent samples.

To investigate the impact of AnkB absence on lens fiber protrusions/paddles and the distribution of various membrane-associated proteins, we isolated and immunostained fibers derived from P16 lenses (from inner cortical region) for AnkB, β-spectrin, periaxin, and NrCAM, and stained for F-actin. Single fibers were imaged using a Zeiss 880 Airyscan super resolution confocal microscope. This analysis revealed that cortical fibers isolated from AnkB cKO lenses were significantly shorter compared to controls (**Fig. 5B**) and displayed disruptions in and loss of lateral membrane protrusions/paddles (**Fig. 5B**, arrows), along with decreased staining for F-actin, β-spectrin, periaxin, and NrCAM relative to control fibers. AnkB revealed its distribution to the protrusions and paddles of fiber cells very similar to NrCAM, β-spectrin, and β-actin in control lenses (**Fig. 5B**). Additionally, single fiber cells derived from P16 AnkB cKO mouse lenses showed a dramatic disruption in the clustering of connexin-50 (Cx-50) and β-dystroglycan at the broad side of the lateral membrane compared to control cells (**Fig. 6**; arrows). Interestingly, despite decreased levels of periaxin, NrCAM, N-Cadherin, and β-spectrin, analysis of lens membrane-enriched fractions from AnkB cKO mice indicated significant increases in the levels of Cx-50 and β-dystroglycan compared to controls (**Fig. 6**), while the levels of dystrophins (DP71 & DP40) decreased significantly in the absence of AnkB, relative to controls (**Fig. 4**).

**Figure. 6.**
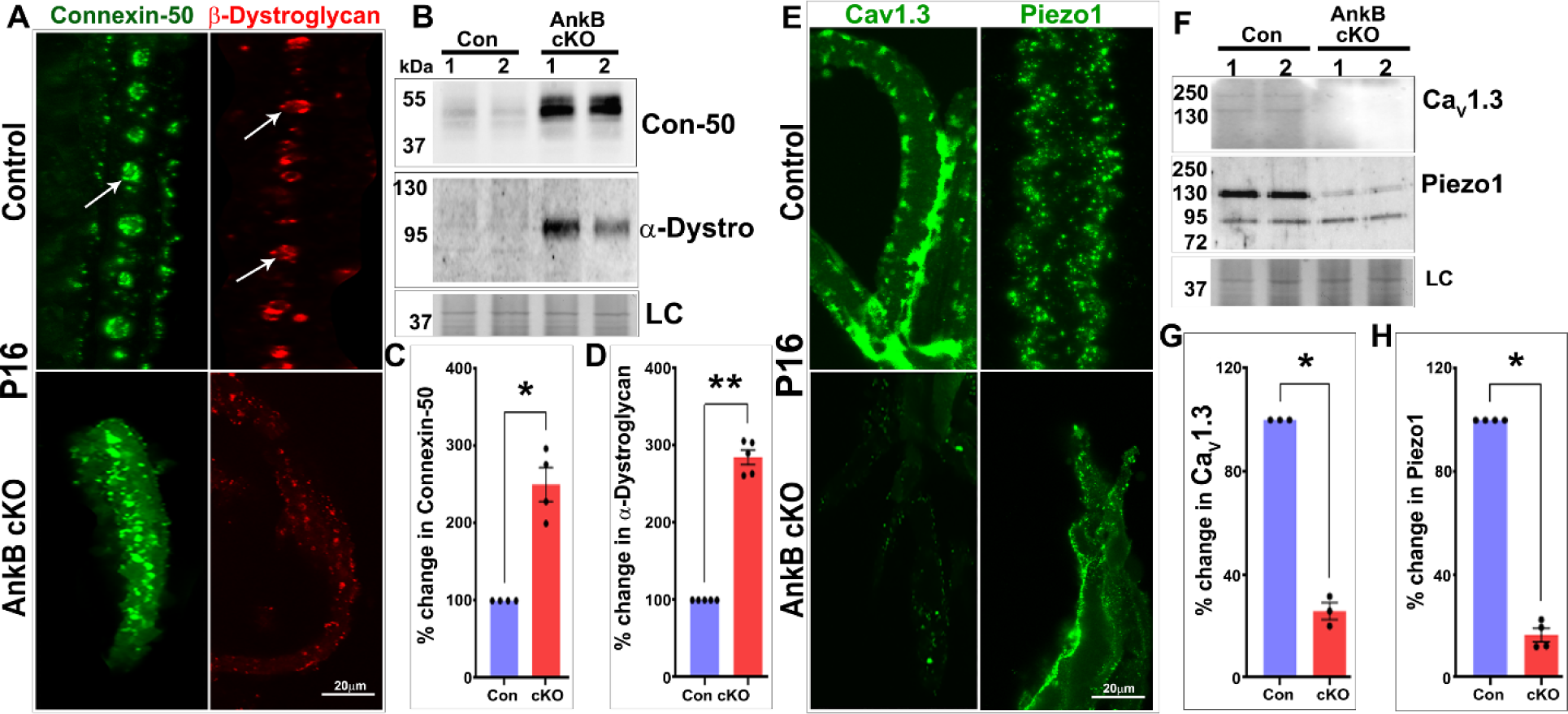
Ankyrin-B deficiency disrupts membrane organization of key channel proteins in lens fibers. Confocal microscopy images of single lens fibers from P16 control (AnkB floxed) and AnkB cKO mice were stained to assess the membrane organization of connexin-50, β-dystroglycan, Ca_V_ 1.3 and Piezo-1 proteins. In control mice, connexin-50 and β-dystroglycan exhibited a distinct round ball-like distribution at the center of the long/broad arm of fiber cells (A, arrows). However, in AnkB cKO mice, the membrane organization of connexin-50 and β-dystroglycan was completely disrupted, showing a scattered pattern (A). Immunoblot analysis of membrane enriched fractions from P16 AnkB cKO lenses revealed a significant increase in levels of connexin-50 and β-dystroglycan compared to controls (B, C and D). Interestingly, the distribution pattern of Ca_V_ 1.3 and piezo-1 channel proteins differed from connexin-50 and β-dystroglycan, localizing to lateral membrane protrusions and paddles in control mice (E). However, in AnkB cKO mice, their distribution was disrupted, and levels in the membrane-enriched fractions were significantly decreased compared to control lenses (panels E, F, and G). LC: Loading control, Con-50: connexin-50, α-Dystro: α-dystroglycan. * P<0.05, Bars represent magnification scale. Lanes 1 and 2 represent two independent samples. Images are representative of analyses from 3 to 4 independent samples.

Furthermore, the distribution of Piezo1 and Ca_v_1.3 channels was noted to be disrupted and decreased at the short sides of the fiber cell lateral membrane in AnkB cKO mouse lens fibers compared to control fibers (**Fig. 6**). These findings were associated with a significant decrease in the levels of Piezo1 and Ca_v_1.3 proteins in the membrane fraction of AnkB cKO lenses compared to control samples (**Fig. 6**). Collectively, these findings suggest that the absence of AnkB disrupts the spectrin/actin-dependent cell adhesion of lens fibers and the membrane organization of NrCAM, Cx-50, calcium channel (Ca_v_1.3), and pressure-sensitive Piezo1 channel in lens fibers.

### Loss of lens stiffness in AnkB cKO mice

Fiber cells peeled from cryopreserved AnkB cKO lens sections (from cortical region) appeared shorter and more fragile compared to those from control lenses (Supplemental **Fig. S8**). Notably, while fiber cells from control lenses curled into worm-like forms upon peeling, those from AnkB mouse lenses remained static, suggesting potential alterations in fiber cell mechanical properties in the absence of AnkB (Supplemental **Fig. S8**). Moreover, in line with previous studies demonstrating reduced lens stiffness in AnkB haploinsufficient, dystrophin-deficient, and periaxin-null mice (Karnam et al., 2021, Maddala et al., 2016), we found a significant decrease (approximately 35%) in lens stiffness (Young’s modulus determined using a microstrain analyzer) in P14 AnkB cKO mice compared to corresponding controls (n=12) (**Fig. 7**). These findings consistently imply AnkB’s involvement in the maintaining lens mechanical properties, potentially through its influence on the spectrin-actin cytoskeleton, cell adhesive interactions, membrane organization, and channel protein activity.

**Figure 7.**
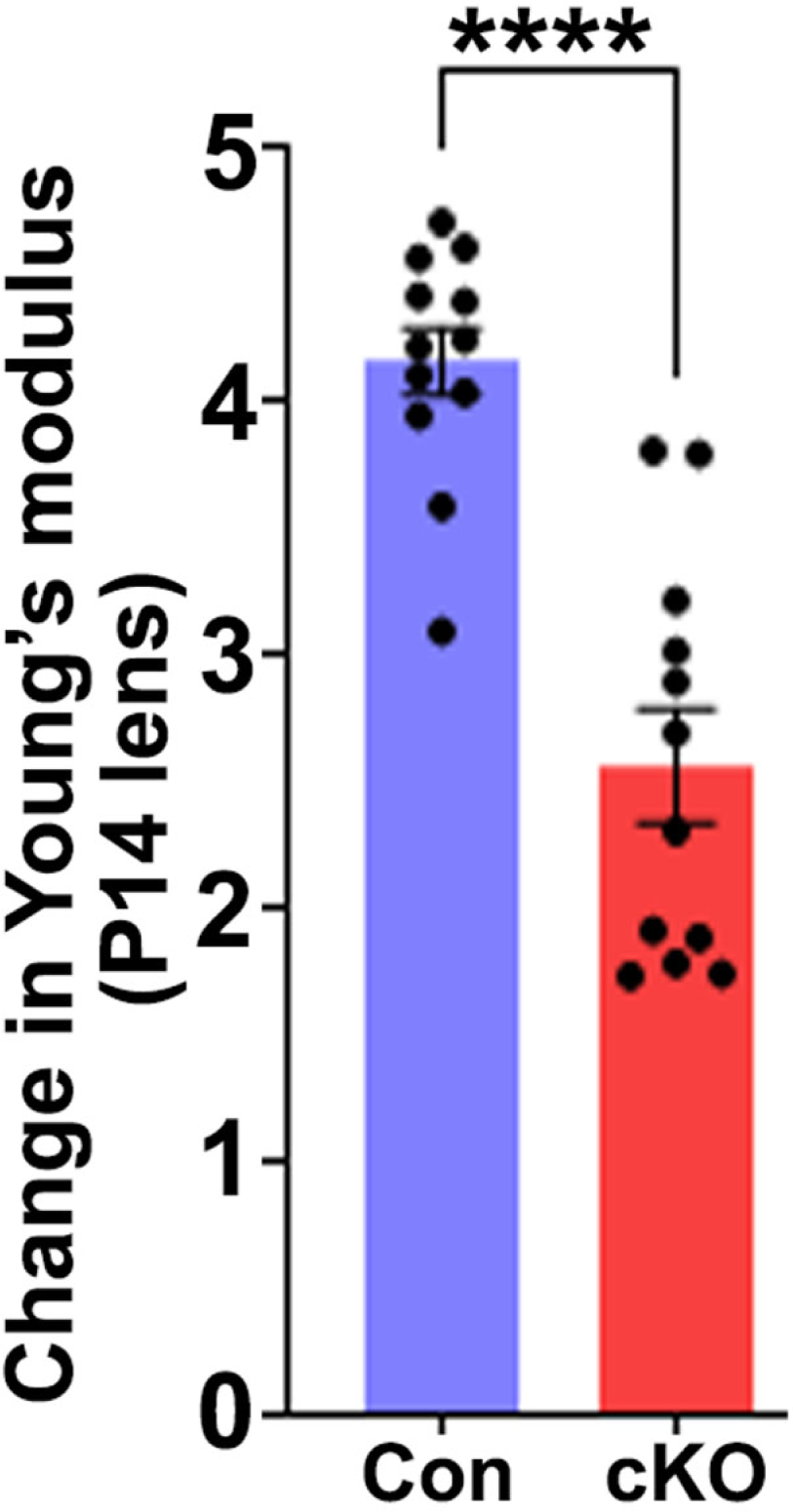
Ankyrin-B deficiency disrupts lens mechanical properties. To assess the effects of AnkB deficiency on lens stiffness, lenses from both control (AnkB floxed) and AnkB cKO mice at postnatal day 14 (P14) were analyzed using a microstrain analyzer. Although lenses from both groups appeared transparent, those from the AnkB cKO mice exhibited a significant decrease (by ∼35%, * P<0.05) in Young’s modulus (a measure of stiffness) compared to control lenses. These findings highlight the role of AnkB in modulating the mechanical properties of the lens.

## DISCUSSION

The clarity and accommodation of the ocular lens rely heavily on its distinctive cellular architecture, compaction, microcirculation, and tensile properties (Quinlan and Clark, 2022, Hejtmancik et al., 2015, Mathias et al., 2010). Our study elucidates the crucial role of ankyrin-B (AnkB), a membrane scaffolding protein, in lens growth, clarity, and mechanical attributes by characterizing lenses from the AnkB cKO mouse model. While membrane protein organization and activities necessary for organs and tissue development and growth are regulated by scaffolding proteins and membrane-tethered spectrin-actin cytoskeletal networks (Bennett and Baines, 2001), our understanding of the specific role of AnkB, an abundantly expressed membrane scaffolding protein, in lens growth and function remains limited (More et al., 2001, Maddala et al., 2016). This study unveils the conditional deficiency of AnkB affecting lens growth and function without disrupting its development and differentiation, primarily by impairing fiber cell morphology, adhesion, membrane protein stability and organization, as well as tensile properties. **Figure 8** schematically illustrates the influence of AnkB on various aspects of the lens, including growth, function, and fiber cell characteristics.

**Figure 8.**
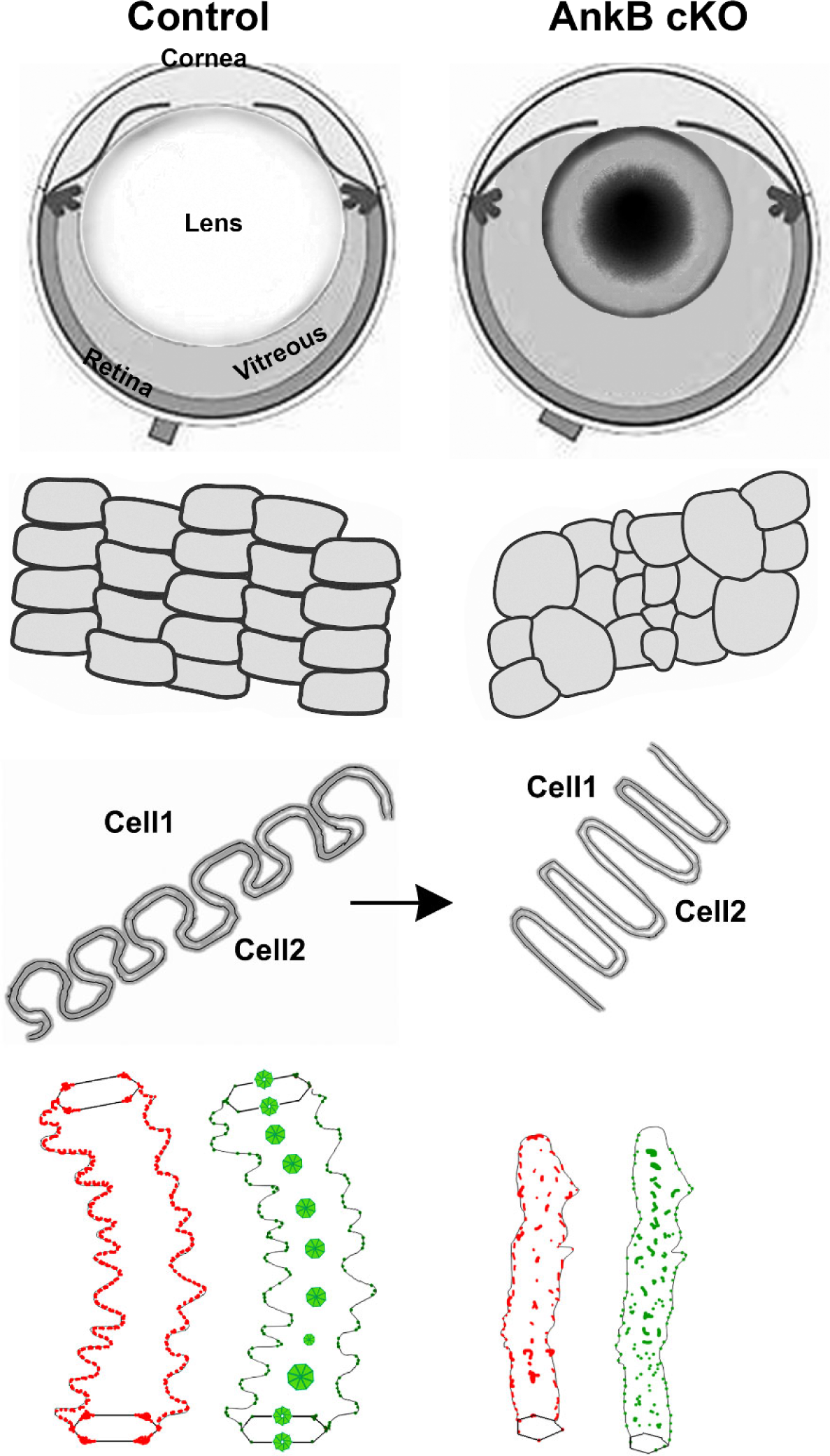
Schematic illustration of the impact of Ankyrin-B deficiency on lens growth, function, and fiber cell architecture.

The differentiation of cuboidal epithelial cells at the equatorial regions of the lens into long fiber cells, which occurs continuously throughout life, underpins lens secondary fiber differentiation (Hejtmancik et al., 2015, Cvekl and Ashery-Padan, 2014). As new fibers overlay the older ones concentrically, akin to the layers of an onion, the older fibers migrate towards the lens center. Outer and inner cortical fibers acquire hexagonal symmetry and radial arrangement (Cheng et al., 2017, Quinlan and Clark, 2022). Notably, as fibers differentiate and migrate inward, they develop elaborate lateral membrane ball- and-socket interdigitations in the cortical region, enhancing compaction (Taylor et al., 1996, Quinlan and Clark, 2022). These fibers express high levels of membrane and membrane-associated proteins such as AnkB, NrCAM, connexins, and aquaporin-0 (Al-Ghoul et al., 2001, Blankenship et al., 2007, Mathias et al., 2010, Li et al., 2022, More et al., 2001, Gu et al., 2019). Additionally, fibers contain the spectrin-actin cytoskeleton tethered to the membrane (Cheng et al., 2017). Fiber cell shape, adhesion, compaction and microcirculation rely on the organization and activities of various membrane proteins, including cell adhesion, transport, channel, and signaling proteins (Cheng et al., 2017, Mathias et al., 2010, Li et al., 2022, Maddala et al., 2013, Gu et al., 2019, Shiels and Bassnett, 1996, Wang et al., 2016, Li et al., 2023). Scaffolding proteins linking the spectrin-actin cytoskeleton to membrane proteins are essential for their stability and activity (Bennett and Lorenzo, 2016). Here, we focus on elucidating the role of AnkB, which is abundantly expressed in lens fibers, in lens cytoarchitecture, growth and function.

Ankyrin-B gene variants are associated with an increased risk of various diseases, including cardiac, neurological, and metabolic defects (Stevens and Rasband, 2021, Mohler and Bennett, 2005, Lorenzo et al., 2015), suggesting potential implications for lens physiology and function. Previous studies on AnkB null mice have reported disorganized fibers, but due to their early demise, the precise role of AnkB in lens growth and function remains unclear (More et al., 2001). The AnkB cKO mouse model developed in this study allows analysis and characterization of lens abnormalities at and post P14 stages, providing insights distinct from AnkB null mice (More et al., 2001). Additionally, the overt phenotype observed in AnkB cKO mice was not evident in AnkB haploinsufficient mice (Maddala et al., 2016), further underscoring the significance of the AnkB cKO model for understanding AnkB’s role in lens biology.

One intriguing finding in AnkB cKO mice is the discrete decrease in lens size and weight starting at post P14, accompanied by intense opacity in the nucleus. The rapid reduction in lens size and weight at and post P16, associated with complete opacification, highlights the critical period between P14 and P16 for AnkB-mediated lens growth and function. The mechanisms underlying this phenomenon warrant further investigation, particularly regarding abnormalities in fiber cell lateral membrane interdigitations under AnkB deficiency (Blankenship et al., 2007). Notably, while wet weight robustly increased from P14 to P21 in normal lenses, this trend was absent in AnkB cKO lenses. Although apoptosis was not observed, extensive membrane folding, fragmentation, and protein degradation, along with impaired lateral membrane interdigitations and cell adhesion, were evident in AnkB cKO lenses, indicating abnormalities in membrane characteristics regulated by AnkB. Furthermore, AnkB deficiency appears to disrupt epithelial phenotype based on αSMA expression and decreased E-cadherin immunostaining, suggesting potential secondary effects on epithelial cells due to AnkB absence. Future studies are needed to elucidate the direct impact of AnkB deficiency on epithelial phenotype.

Our study provides compelling evidence for the importance of AnkB in regulating fiber cell morphology, adhesion, and membrane skeletal organization in lens growth and function. We observed increased instability of membrane and membrane-associated proteins, including NrCAM, spectrin, N-cadherin, periaxin, aquaporin-0, and dystrophins in AnkB ckO lenses, indicating a crucial role for AnkB in maintaining their stability and organization. Colocalization analyses revealed AnkB’s close interaction with various membrane and membrane-associated proteins including NrCAM, β-spectrin and periaxin, highlighting its influence on their stability and organization in lens fibers. Despite disruption in the membrane domain organization of Cx-50 and β-dystroglycan in AnkB cKO lenses, their elevated protein levels suggest a compensatory response to AnkB deficiency. Similar to AnkB, Cx-50 and aquaporin-0 deficiencies impair the development of ball-and-socket interdigitations in lens fibers (Wang et al., 2016, Gu et al., 2019), emphasizing the collective role of membrane scaffolding, cytoskeletal, and channel proteins in maintaining fiber cell lateral membrane interdigitations. Moreover, the decreased lens stiffness observed in AnkB cKO mice underscores AnkB’s role in maintaining mechanical properties essential for lens accommodation, consistent with previous findings in AnkB haploinsufficient mice (Maddala et al., 2016).

In conclusion, our study utilizing an AnkB cKO mouse model highlights the indispensable role of this membrane scaffolding protein in maintaining fiber cell architecture, adhesion, compaction, membrane organization, and mechanical properties, thereby regulating lens growth and function. The absence of AnkB exerts detrimental effects on this vital visual organ.

## MATERIALS AND METHODS

### Mice

All experiments using mice were carried out in accordance with the recommendations of the Guide for the Care and Use of Laboratory Animals of the National Institutes of Health and the Association for Research in Vision and Ophthalmology. The protocol (A213-19-10) was approved by the Institutional Animal Care and Use Committee (IACUC) of the Duke University School of Medicine. Mice were maintained in a pathogen-free vivarium under a 12-hour dark and light cycle with ad libitum food and water.

### Generation of Ankyrin-B conditional knockout (AnkB cKO) mice

To investigate the role of AnkB in lens growth and function, we generated AnkB cKO mice by mating AnkB floxed mice with loxP sites flanking exon 24 of the *ANK2* gene (maintained on a C57BL/6J background) obtained from Peter Mohler (Smith et al., 2015), The Ohio State University Wexner Medical Center with lens-specific Cre recombinase expressing transgenic mice (Le-Cre mice) as we described earlier (Maddala et al., 2015). The Le-Cre transgenic mice utilized in this study express Cre recombinase at embryonic day 8.75 under the control of a Pax6 P0 enhancer/promoter. Cre expression occurs in the lens epithelium and fiber cells, and other surface ectoderm-derived eye structures(Ashery-Padan et al., 2000). For comparison analysis, littermate AnkB floxed controls lacking the Cre transgene alongside AnkB cKO mice were used. AnkB cKO mice, bred for 10 or more generations on a C57BL/6J genetic background, were compared with these controls. To confirm the presence of AnkB floxed alleles and the Cre transgene in the progeny, tail DNA was subjected to PCR analysis using the following oligonucleotide primers:

AnkB: forward/reverse primers:

GCAGTCTCAACACAACTAAGCCATCCTTTT GCTGAGGAGGTAGACAAGAACCTTTTTGTG

Cre forward /reverse primers:

GCATTACCGGTCGATGCAACGAGTGATGAG GAGTGAACGAACCTGGTCGAAATCAGTGCG

### Tissue fixation and sectioning

#### Paraffin sections

Whole eyes from P1 and various other ages of wild type (WT), AnkB cKO and littermate AnkB floxed (control) mice underwent fixation in 10% buffered formalin for 48 hrs. To ensure proper fixation, a small incision was mode at the optic stalk for optimal permeation of the fixative. Following fixation, the specimens were dehydrated and embedded in paraffin. Subsequently, 5µm thick sections were cut using a Leica Leitz 1512 microtome, either in the sagittal or equatorial plane, and stored at room temperature (RT) until further processing.

#### Cryosections

Whole eyes from WT, AnkB cKO, and AnkB floxed mice at various developmental stages (P1, 10, 14, 16, 18 and 21 day) were fixed in 4% buffered paraformaldehyde for 24 hrs at 4 °C. The fixed tissues were then subsequently transferred into 5% and 30% sucrose solutions in phosphate-buffered saline (PBS) over consecutive days. After embedding in optimal cutting temperature media (Tissue-Tek, Torrance, CA, USA), the samples were sectioned into 10 μm-thick sagittal sections using a Microm HM550 Cryostat (GMI, Ramsey, MN, USA) and stored at − 80 °C until further use.

#### Lens single cell isolation

At postnatal day 16 (P16), lenses were carefully dissected from both AnkB floxed and AnkB cKO mice and fixed in 4% paraformaldehyde (pH7.4) for 2 hrs at room temperature, with extra precautions taken to avoid rupturing the AnkB cKO lenses. The lens capsule was then carefully peeled off, and the fixed lenses were quartered in the anterior to posterior pole direction using a sharp razor blade, followed by an additional 2 hr fixation period. Subsequently, the lens nuclear regions were removed and discarded using tweezers, while the remaining fiber layers were peeled and collected at different depths according to the method described by Chen et. al. (Cheng et al., 2018).

### Immunofluorescence staining and imaging

#### Paraffin sections

Paraffin-embedded sections (5μm thick) in either the equatorial or sagittal plane underwent de-paraffinization and rehydration using xylene and absolute ethyl alcohol, as previously described (Maddala et al., 2011). Subsequently, specimens were subjected to antigen retrieval by immersion in preheated antigen retrieval solution (0.1 M Na Citrate buffer pH 6.0) at 100 °C for 20 min. Following rinsing, sections were blocked for 10 min using medical background Sniper reducing solution (BiocareMedical) in a humidified chamber. Sections were then treated with primary antibodies, including anti-AnkB antibody (rabbit polyclonal antibodies, generously provided by Vann Bennett), in conjunction with either anti-NrCAM, anti-β-spectrin, anti-periaxin, β-Actin, connexin-50 or β-dystroglycan antibodies for colocalization analysis. All antibodies (Supplemental **Table S2**) were used at 1:200 dilution in 1% fatty acid-free BSA in Tris buffer saline and incubated overnight at 4 °C in a humidified chamber. Subsequently, slides were washed and incubated in the dark for 2 hrs at RT with the respective Alexa fluor 488- and 568-conjugated secondary antibodies (Invitrogen; at 1:200 dilution). After further washing, slides were mounted using Vecta mount (Vector Laboratories), with edges sealed using nail enamel prior to image capture.

#### Cryosections

Air-dried tissue cryosections underwent treatment with Image-iT FX signal enhancer (Invitrogen, Eugene, OR, USA) followed by blocked in blocking buffer (5% globulin-free bovine serum albumin and 5% filtered goat serum in 0.3% Triton X-100 in PBS) for 30 min each. Tissue sections were then incubated with tetra rhodamine isothiocyanate–conjugated phalloidin (TRITC; Sigma Aldrich) at a 1:500 dilution for 2 hrs at RT for F-actin staining. Following washing, slides were mounted using Vecta mount, and edges were sealed with nail enamel.

#### Single fibers

Fiber stacks peeled from cryopreserved lens sections were blocked with blocking buffer (5% BSA (fatty acid free), 5% normal goat serum in 0.3% triton X 100 in1X PBS) for 1hr at RT. They were then incubated with respective primary antibodies (Supplemental **Table. S2**) for 2 hrs at RT, followed by washing with wash buffer (0.3% triton X 100 in1X PBS) and incubation with respective Alexa Fluor 488- or 568-tagged secondary antibodies for 1hr at RT. Single layers were carefully peeled from the immunostained fiber stacks and mounted under glass coverslips. Images were captured using a Nikon Eclipse 90i confocal laser scanning microscope (100X 1.40 oil). Immunostaining was repeated on four independent lenses from Floxed and cKO lenses, with representative data shown.

#### Z stack imaging

Z stack images were captured at 0.5μm interval using a Nikon Eclipse 90i confocal laser scanning microscope (100X 1.40 oil). We obtained 15 optical sections for each sample. All representative immunofluorescence data reported in this study were based on maximum intensity projections of the Z stacks. A minimum of four tissue sections derived from four independent specimens were used as biological replicates.

#### Z imaging and 3D reconstruction

For co-localization measurements and 3D reconstruction, immunostained sections (paraffin) were imaged using a Zeiss 780 airyscan inverted confocal microscope equipped with Argon/2 and 561nm diode lasers. We utilized a 100X 1.40 NA oil objective lens with a zoom factor of 2. Images were acquired using Zeiss 2.3 imaging software. Z-stack images were captured at 0.3μm intervals, resulting in 12 optical sections per sample. Volocity 6.3.1 software (Perkin Elmer) was employed for processing the captured Z-stacks. Coste’s Pearson’s correlation coefficient (CPC) for AnkB co-localization with other proteins was analyzed using the raw images. Deconvoluted images were utilized for 3D reconstruction, processed using Adobe Photoshop version CS4. All representative immunofluorescence data reported in this study were based on a minimum of 3 to 6 tissue sections derived from three independent specimens per group.

### Tissue clearing and Light sheet microscopy

P16 lenses from AnkB cKO and AnkB floxed control mice were utilized for tissue clearing and subsequent light sheet microscopy. Initially, lenses were fixed in 4% paraformaldehyde for 24 hrs. Subsequently, small pores were created in the lens capsule, and the lenses were fixed for an additional 24 hrs. The lenses were then subjected to a series of methanol concentrations (25%; 50%; 75% and 100%), with each concentration applied for one hr, followed by a reversal of the order of methanol concentrations every 15 min. Finally, the lenses were stored in PBS before treatment with CUBIC reagents. All incubations with CUBIC reagents were carried out at 37 ^0^C on a circular rocker. The lenses were initially incubated in CUBIC reagent1 (containing 25g of urea, 80% Quadrol (N, N, N, N-Tetrakis(2-Hydroxypropyl) ethylenediamine, (cat no: 122262, Millipore Sigma) in H_2_O, 15 ml TritonX100), at a 50% concentration in PBS for one day, followed by three days at 100% concentration, with the reagent changed every 24 hrs. Subsequently, the lenses underwent rinsing with 1X PBS for 3 times for 30 mins each, followed by permeabilization and washing with buffer (1X PBS with 2% TritonX100) for 3 hrs. The lenses were then blocked over night at 4^0^C while rocking, in a solution containing 10% FBS and 2% TritonX100 in PBS. Following blocking, the lenses were incubated with primary antibody against Aquaporin 0 (AQP0) at a 1:1000 dilution in blocking buffer for 48 hrs. After washing with PBS containing 2% TritonX100, the lenses were incubated with a secondary antibody (Alexa Fluor 488) at 1:2500 dilution in blocking buffer overnight. Subsequent to another round of washing with the buffer for three one-hr intervals at RT, the lenses were embedded in 1% low melting agarose. The agarose blocks were then transferred into CUBIC reagent 2 (comprising 25g of urea, 50g sucrose, and 10 ml of Triethanolamine; cat. No: 1371481000, Millipore Sigma) at a 50% concentration for 24 hrs, followed by three days at 100% concentration, with daily replacement of the reagent until complete tissue clearing was achieved. Finally, the agarose-embedded specimens were imaged on a light-sheet microscope (Ultramicroscope II, LaVision BioTec) equipped with an sCMOS camera (Andor Neo) and a 2X /0.5 objective lens with a 6mm working distance dipping cap.

### Transmission electron microscopy

For transmission electron microscopy (TEM)-based histological analysis, lenses from P14 and P16-day-old AnkB floxed and AnkB cKO mice were fixed in 10% buffered formalin with 0.25% glutaraldehyde for 48 hrs. The fixed tissue specimens were subsequently treated with 1% osmium tetraoxide in 0.1 M sodium cacodylate buffer (pH 7.2) at 4 °C for 1hr. After rinsing with water and ethanol, the tissue specimens were treated with 2% uranyl acetate for 1 hr at RT. The samples were then dehydrated through a series of ethanol grades (30% to 100%) before being infiltrated with a 1:1 mixture of propylene oxide and Spurr resin (SPI supplies, West Chester, PA), followed by embedding in pure Spurr resin at 65 °C overnight. The tissue specimens were sectioned (0.7 µ) in the equatorial plane using a Reichert Ultra-cut microtome (Leica), followed by production of ultra-thin sections (65-75 nm). Tissue sections were treated with 2% uranyl acetate, rinsed with 25% ethanol and water, and treated with a 3.5% lead citrate solution. Images were captured at 8000x using a Jeol JEM-1400 equipped with an Orius CCD digital camera at 60kV (JEOL, Tokyo, Japan).

### Scanning electron microscopy

For scanning electron microscopy (SEM) imaging, lenses from P14 and P16-day-old AnkB floxed and AnkB cKO mice were fixed in 2% buffered paraformaldehyde with 2.5% glutaraldehyde in 0.1M sodium cacodylate buffer for 1 hr. The tissue specimens were then washed three times for 5 min each with 0.1 M cacodylate buffer and post fixed with 1% osmium tetroxide (OsO_4_)/0.8% potassium ferrocyanide (K_4_Fe (CN)_6_) in 0.1M cacodylate, pH 7.4 for 30 min. After further washing with 0.1 M cacodylate buffer, followed by two washes with distilled water, dehydration was carried out in 95% ethyl alcohol initially for 1 min followed by three washes of 5 min each in 100% ethanol. Specimens were then allowed to dehydrate overnight in a desiccator with activated desiccant. The following day, specimens were secured on aluminum stubs, carefully fractured, sputter-coated with gold particles, and imaged using a Hitachi TM3030Plus tabletop scanning electron microscope equipped with secondary electron and backscattered electron detectors.

### Immunoprecipitation Analysis

To elucidate the interactome of AnkB, immunoprecipitation (IP) assays were conducted using a monoclonal antibody against AnkB (Invitrogen) in conjunction with Dynabeads Protein-G (Invitrogen), following previously established protocol (Maddala et al., 2011). Briefly, mouse lenses, obtained as pooled samples from P21 to P30 mice, were homogenized (2% w/v) in 1× IP buffer (10X IP Buffer, Cat. No. I5779, Sigma-aldrich.com) supplemented with protease and phosphatase inhibitors (protease inhibitor tablet, Cat. No 11836170001, and PhosStop tablet, Cat. No. 04906837001, Roche Diagnostics. IN). Homogenization was performed on ice using a Dounce glass homogenizer with 10 upward and 10 downward strokes. The homogenates were then centrifuged at 800×g for 10 min at 4 °C, and the resulting supernatants were collected. Protein concentrations were determined using the Bio-Rad dye reagent prior to use in IP reactions. Immunoprecipitation assays were conducted both for AnkB IP and IgG control under identical conditions. AnkB immunocomplexes were suspended in 2× Laemmli sample buffer and boiled for 5 min. The samples were then resolved by gradient SDS-PAGE (4-20%; Criterion XT precast gel) and stained with Gelcode® Blue stain reagent. Following destaining with Milli Q pure water, protein bands of interest were excised from the gel and subjected to in-gel tryptic digestion using the In-Gel Tryptic digestion kit (Pierce), following the manufacturer’s instructions. This digestion process involved both the reduction and alkylation of protein samples, which subsequently identified by mass spectrometry.

### Mass spectrometry

Peptides obtained from in-gel-trypsin-digests, as previously described (Maddala et al., 2011) were subjected to analysis a nanoAcquity UPLC system coupled to a Synapt G2 HDMS mass spectrometer (Waters Corp, Milford, MA). For each sample, we employed a data-dependent analysis (DDA) approach. This involved an initial 0.8 s MS scan followed by MS/MS acquisition of the top three ions with a charge greater than one. MS/MS scans for each ion were performed using an isolation window of ∼3 Da. To enhance data quality, a maximum acquisition time of 2 s per precursor was imposed. Additionally, dynamic exclusion was applied for 120 s within 1.2 Da to minimize redundant spectra acquisition. DDA-generated data were processed using Protein Lynx Global Server 2.4 (Waters Corporation). Processed data were then searched against the NCBI mouse database using Mascot server 2.2. Identification of proteins was based on the detection of at least two peptides for each protein. Additionally, a confidence interval percentage (CI %) of over 99.9% was required for inclusion, corresponding to a false discovery rate of 0.1%.

### Immunoblotting

Lens tissue from postnatal day 12, 14, 16, and 18 AnkB cKO and floxed control mice was pooled and homogenized using a Dounce glass homogenizer in a cold (4°C) hypotonic buffer. The buffer composition included 10 mM Tris buffer pH 7.4, 0.2 mM MgCl_2_, 5 mM N-ethylmaleimide, 2.0 mM Na_3_VO_4_, 10 mM NaF, 60 µM phenyl methyl sulfonyl fluoride (PMSF), 0.4 mM iodoacetamide, protease inhibitor cocktail (complete, Mini, EDTA-free), along with PhosSTOP phosphatase inhibitor. The homogenates were centrifuged at 800×g for 10 min at 4°C, followed by a second centrifugation at 100,000×g for one hr at 4°C. The supernatant obtained from the second centrifugation was designed as the cytosolic fraction. The insoluble pellets were re-suspended in hypotonic buffer, washed twice with same buffer, and the resulting membrane-enriched protein pellets were suspended in urea sample buffer containing 8 M urea, 20 mM Tris, 23 mM glycine, 10 mM dithiothreitol (DTT), and saturated sucrose along with protease and phosphatase inhibitors. Protein concentrations in both the cytosolic and membrane-enriched fractions were estimated using the Micro BCA™ Protein Assay Kit (Cat no. 23235, Thermo Scientific). Equal amounts of protein samples were separated by SDS-PAGE ((4-20%; Criterion XT precast gel) with 1× MOPS SDS gel running buffer (NP0001, Invitrogen, Life Technologies Corp). Following electrophoresis, gels were stained with Gelcode® Blue stain reagent and imaged using the ChemiDoc™ Touch Imaging System.

For immunoblotting, membrane enriched fractions were resolved on either 8 or 10% SDS-PAGE gels based on the molecular mass of the protein of interest. Proteins were then transferred electrophoretically onto nitrocellulose membrane. Membranes were blocked with 5% nonfat dry milk in 0.1% Tween-20 containing Tris-buffered saline and then incubated with primary antibodies (AnkB, NrCAM, Periaxin, Dystrophin, β-Spectrin, αN-catenin, aquaporin-0, N-Cadherin, connexin-50, α-dystroglycan), piezo-1, and Ca_v_1.3 at a 1: 1000 dilution over night at 4 ^O^C (**Table S2**). After rinsing, membranes were incubated with second antibodies for 2 hrs at RT. Immunoblots were developed using enhanced chemiluminescence (ECL), and images were captured with the ChemiDoc™ Touch Imaging System (Bio-Rad laboratories, Hercules, CA).

### TUNEL assay

To assess and compare apoptotic cell death between AnkB cKO and AnkB floxed control mice, lens cryosections obtained from postnatal days, 10, 14, 16 were subjected to in-situ Terminal deoxynucleotidyl Transferase dUTP Nick End Labeling (TUNEL) using the ApopTag Plus Fluorescein Kit (EMD Millipore, Burlington, MA, USA), following previously established protocols (Maddala et al., 2015). Imaging was performed using a Nikon Eclipse 90i confocal microscope. Representative data in this study were based on the analysis of a minimum of four tissue sections derived from four independent specimens.

### Biomechanical tension compression analysis

The biomechanical stiffness of P14 day-old mouse lenses was analyzed using an RSA III micro strain analyzer (TA Instruments, New Castle, DE), as previously described (Maddala et al., 2016). Lenses from AnkB cKO and AnkB floxed control mice were dissected and placed into organ culture media comprising Dulbecco’s Modified Eagle Medium (Low Glucose DMEM, 298 mOsm), supplemented with penicillin (100 U/ml) and streptomycin (100 mg/ml). Dissected lenses were temporarily stored in a 37°C incubator under 5% CO_2,_ with ruptured and procedurally injured lenses being discarded. Briefly, compression analyses were conducted between two 8mm plates, attached to parallel plate tools and mounted on actuator shafts. Measurements were performed at an ambient temperature of 22-25°C, with lenses submerged in organ culture media. The lens-containing plate was placed on the lower parallel plate, ensuring central alignment, before lowering the upper plate to make contact with the lens specimen. The lens was then strained at a constant rate of 0.05 mm/second for 35 seconds until rupture (typically occurring around 30 seconds for adult lenses). Data were plotted in real time, and TA Orchestrator software facilitated data acquisition. Strain was calculated by measuring changes in applied force divided by the area. Slopes before and after ruptures were determined, with slopes just before rupture plotted using Microsoft excel, as previously described (Maddala et al., 2016).

### Statistical analysis

We conducted a Student’s t-test to assess the significance of differences between the AnkB cKO and AnkB floxed littermate lens specimens. All the values are represented as Mean ± Standard Error of the Mean (SEM). A difference with a p-value less than 0.05 was deemed statistically significant.

## Acknowledgements

We are grateful to Peter Mohler for generously sharing the Ankyrin-B floxed mice used in this study. Our special thanks to Vann Bennett for sharing the AnkB null mice, AnkB polyclonal antibody, and for his unconditional advice during this study. We thank Mark Walters for his help in microstrain analysis, Michelle Gignac for her assistance in scanning electron microscopy, and Ying Hao for her technical support in transmission electron microscopy. Furthermore, we acknowledge with appreciation the support and assistance provided by the Duke University core facilities, particularly the Shared Materials Instrumentation and Light Microscopy units.

## Data Availability

All data referenced in this manuscript, including supplementary materials, are fully disclosed. Additional data can be made available upon request to the corresponding author.

## Author Contributions

Conceptualization: R.M., P.V.R.; Methodology; R.M., A.A., N.P.S.; Validation: R.M., N.P.S., P.V.R.; Formal analysis; R.M., N.P.S., P.V.R.; Resources; P.V.R., Data curation; R.M., N.P.S., P.V.R.: Writing: R.M., P.V.R., Reviewing and Editing; R.M., A.A., N.P.S., P.V.R.; Visualization; R.M., P.V.R.; Project administration; R.M., P.V.R.; Funding acquisition: P.V.R.

## Supplemental Material

**Fig. S1A**: **SDS-PAGE analysis of immunoprecipitates from mouse lens homogenates using AnkB monoclonal antibody compared with IgG antibody**. Equal amounts of lens total lysate protein (from pooled P21 and P30 lenses) underwent immunoprecipitation analysis with AnkB and IgG antibodies. Immunoprecipitated proteins separated on SDS-PAGE were stained with Coomassie blue. Arrows indicate prominent proteins.

**Fig. S1B. Immunoblot analysis of immunoprecipitated proteins from mouse lens homogenates using AnkB antibody.** To validate mass spectrometer-identified proteins in AnkB antibody-precipitated fractions, the AnkB antibody-precipitated protein fraction was subjected to immunoblot analyses with specific antibodies.

**Fig. S2. Normal lens development and fiber cell elongation/differentiation in AnkB null and AnkB cKO mice.** Comparison of P1 lens sagittal sections (paraffin embedded) from AnkB null (A) and AnkB cKO (B) mice with respective wild type and AnkB floxed control, stained with hematoxylin and eosin (H&E) or immunostained for AnkB. Similar patterns of development and growth observed in null, cKO, and wild type lenses. Epi: Epithelium. Bars: Magnification scale.

**Fig. S3. Absence of TUNEL positive staining in AnkB cKO lenses.** TUNEL staining of lens cryosections from P10, P14, and P16 AnkB cKO and littermate control (AnkB floxed) mice show no evidence of apoptosis in AnkB cKO mice. Epi: Epithelium. Bars: Magnification scale. Images are representative of analyses from 3 independent samples.

**Fig. S4: Disruption of lens weight increase during P14 to P21 under AnkB absence in mice.** Dramatic increase in lens weight from P14 to P21 observed in control mice, while AnkB cKO mice show discrete lens phenotypes from P16, including cataract and decreased lens weight. **** P<0.0001.

**Fig. S5: AnkB deficiency disrupts E-cadherin distribution and induces *α*-smooth muscle actin expression in lens.** Immunostaining of P14 and P16 AnkB cKO and littermate control (AnkB floxed) lenses reveals thinner epithelium with reduced E-Cadherin staining and induction of αSMA expression in AnkB cKO lenses. Epi: Epithelium. Bars: Magnification scale. Images are representative of analyses from 3 independent samples.

**Fig. S6. iDISCO tissue clearing and light sheet imaging of AnkB cKO mouse lens.** Whole lenses from P16 control and AnkB cKO mice which were immunostained for aquaporin-0 reveal disrupted fiber organization in AnkB cKO lenses compared to controls. Images are representative of analyses from 3 independent samples.

**Fig. S7. Lens soluble protein profile in AnkB cKO mouse lens.** SDS-PAGE (4-20% gradient gel) analysis of soluble lens fractions (100,000xg supernatant) from P12 to P18 control and AnkB cKO mice show decreased levels of crystallins in P18 AnkB cKO lenses.

**Fig. S8. Images of single lens fibers peeled from the AnkB cKO mouse compared to control mice.** Fibers from AnkB cKO lenses (P16) are shorter and fragile, contrasting with long, curling fibers from control (AnkB floxed) lenses. Images are representative of analyses from 3 to 4 independent samples.

**Supplemental material-Movies**: 3D rendering movies showing AnkB co-distribution with β-spectrin, connexin-50, F-actin, and β-dystroglycan in mouse lens fibers (P21 lens cortical region). Click the image to play the movie.

**Table S1:** Identification of Ankyrin-B interacting proteins in mouse lens homogenates based on immunoprecipitation and mass spectrometry analyses.

**Table S2:** Details of primary and secondary antibodies used in immunoblot and immunofluorescence analyses.

